# Developing NMR methods for macromolecular machines: Measurement of residual dipolar couplings to probe dynamic regions of the ribosome

**DOI:** 10.1101/319251

**Authors:** Xiaolin Wang, John P. Kirkpatrick, Hélène M. M. Launay, Alfonso de Simone, Daniel Häussinger, Christopher M. Dobson, Michele Vendruscolo, Lisa D. Cabrita, Christopher A. Waudby, John Christodoulou

## Abstract

We describe an NMR approach based on the measurement of residual dipolar couplings (RDCs) to probe the structural and motional properties of the dynamic regions of the ribosome. Alignment of intact 70S ribosomes in filamentous bacteriophage enabled measurement of RDCs in the stalk protein bL12, and this information was used to refine a 3D structure of its C-terminal domain (CTD). Orientational constraints on the CTD alignment arising from the semiflexible linker sequence were further probed by a paramagnetic alignment strategy, and provided direct experimental validation of a structural ensemble previously derived from SAXS and NMR relaxation measurements. Our results demonstrate the prospect of better characterising dynamical and functional regions of more challenging macromolecular machines and systems, for example ribosome–nascent chain complexes.

## Introduction

In recent years, X-ray crystallography and cryo-electron microscopy (cryo-EM) have elucidated the details of high-resolution structures of ribosomes, revealing intricate mechanistic information about their function during the translation process^1, 2^. In parallel, NMR-based observations of nascent polypeptide chains emerging from the ribosome are providing unique structural and mechanistic insights into co-translational folding processes^3, 4^. In order to develop further solution-state NMR spectroscopy as a technique for structural studies of dynamic regions within large complexes, we have explored the measurement of residual dipolar couplings (RDCs) within intact ribosomes, focusing in particular on the mobile bL12 protein from the GTPase-associated region (GAR) of the prokaryotic 70S ribosome. RDCs have been used to characterise other macromolecular machines and assemblies, including HIV-1 capsid protein, bacterial Enzyme I, and the 20S proteasome^5–7^. These developments are particularly relevant as macromolecular complexes tend to exhibit a wide variety of functional motions that are challenging to characterise by standard methods.

The GAR is a highly conserved region of both prokaryotic and eukaryotic ribosomes, and is involved in the recruitment and stimulation of the GTPase activity of auxiliary factors associated with the key steps of translation^8^. The prokaryotic GAR includes helices 42–44 and 95 of the 23S rRNA, and ribosomal proteins bL10, bL11 and bL12 (Figure 1a). bL12 is unique among the ribosomal proteins in being present in multiple copies – four copies (two dimers) on the E. coli ribosome^9^ – and has a mobile C-terminal domain (CTD) connected to the ribosome-bound N-terminal domain (NTD) via a disordered linker. This flexibility, which has been proposed to facilitate the recruitment of translation factors from the cellular space to the 30S/50S interface^8, 10, 11^, has precluded structure determination by cryo-EM and X-ray crystallography. Solution-state NMR spectroscopy is therefore uniquely suited to the characterisation of this region. Indeed, ^1^H,^15^N-correlation spectra of 70S ribosomes are found to contain resonances arising almost exclusively from the bL12 stalk^12–14^. These previous studies have found that the structure of the CTD on the ribosome is indistinguishable from that in isolation, and that only two of the four copies are mobile on the NMR timescale.

**Figure 1.**
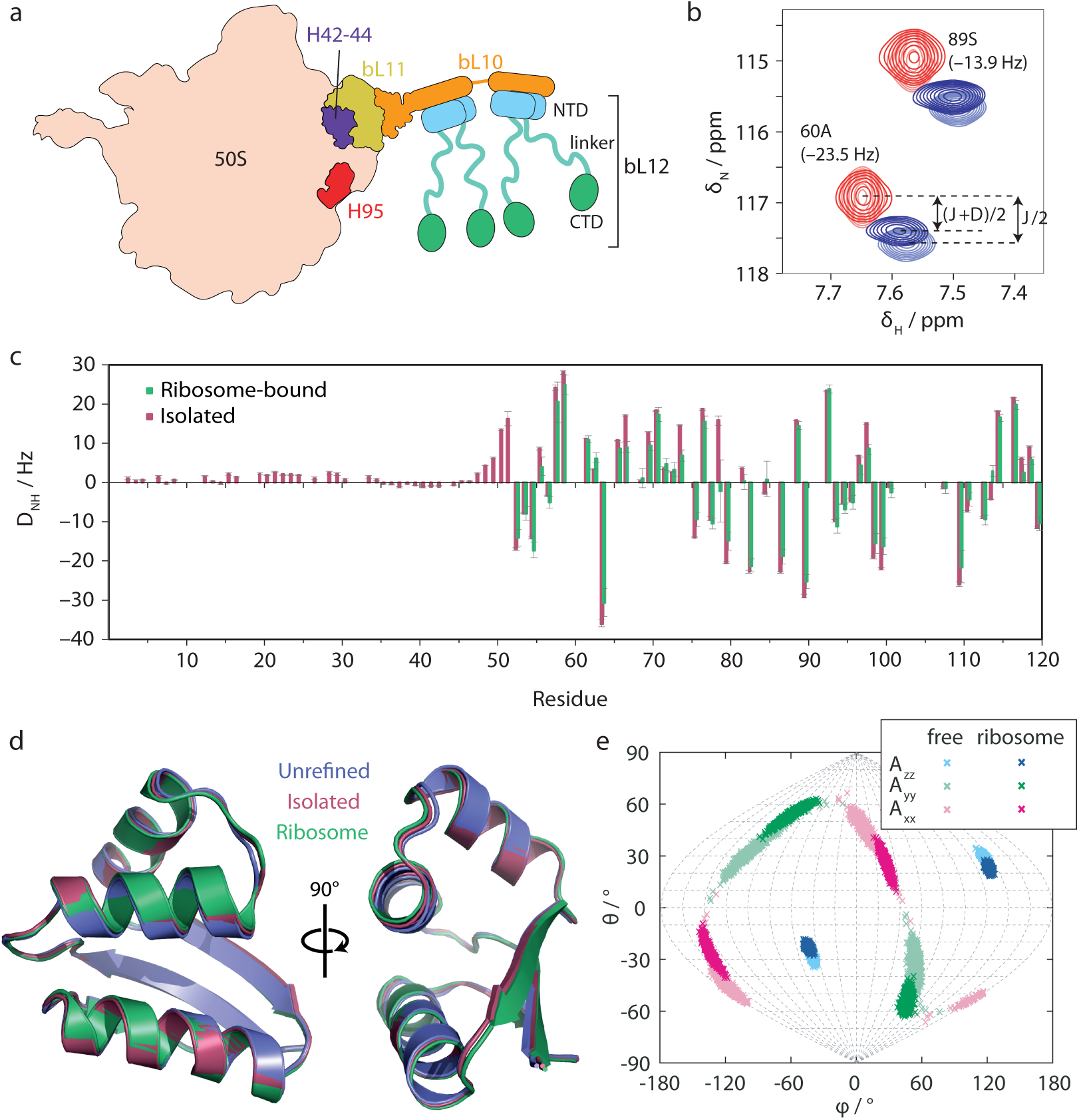
(a) Schematic illustration of the 50S subunit of the bacterial ribosome illustrating the position and composition of the stalk region. (b) Excerpt from ^1^H,^15^N HSQC (red) and TROSY (blue) spectra of the ribosome-bound bL12 CTD in isotropic and aligned conditions (light and dark colouring, respectively), acquired at 700 MHz and 298 K. (c) Amide RDCs measured for ribosome-bound and isolated bL12. (d) Refinement of the bL12 template structure (PDB code:1rqu) using observed RDCs for isolated and ribosome-bound bL12. (e) Sanson-Flamsteed projection of the principal axes of the alignment tensor calculated for free and ribosome-bound bL12.

In this work, we continue our NMR investigations of bL12 by measurement of residual dipolar couplings (RDCs) for this protein in the spectrum of intact 70S ribosomes. RDCs can be measured under weakly anisotropic solution conditions and report on both the local structure of the protein and the overall orientation of domains with respect to a laboratory frame of reference^15, 16^. As such, they are useful sources of distance-independent structural information that are highly complementary to chemical shifts, scalar couplings and NOEs. Anisotropy can be induced by a variety of steric and electrostatic methods including liquid crystals, bicelles, bacteriophage and stretched polyacrylamide gels^17^.

## Results and discussion

We tested the alignment of samples of ribosomes in Pf1 bacteriophage and C_12_E_5_ PEG/hexanol mixtures^18^. The latter induced precipitation and inhomogeneous alignment (methods), but strong alignment was obtained in Pf1 bacteriophage and this system was therefore used for further measurements (Fig. S1). Amide RDCs (*D_NH_*) were determined for both isolated and ribosome-bound bL12 by measurement of ^15^N frequency differences between amide resonances in ^1^H,^15^N HSQC and TROSY spectra (Fig. 1b)^19^. We found that this approach provided the greatest precision in values of RDCs when compared with alternative methods such as the in-phase/anti-phase approach (Fig. S2). Measurements of additional couplings to C’ and C*α* atoms were also explored, but the sensitivity was not sufficient to obtain precise results. The integrity of ribosomal samples was monitored with ^1^H stimulated-echo diffusion measurements, acquired in an interleaved fashion with HSQC and TROSY experiments (Fig. S3)^20^.

Despite the 2.4 MDa molecular-weight of the 70S ribosome, and the low sample concentration (10 *µ*M), RDCs could be obtained for the entire CTD of ribosomal bL12 (bL12_ribo_), with an RMS uncertainty 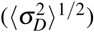 of 1.8 Hz relative to a range of ±30 Hz (Fig. 1c and SI methods). Despite the multiple copies of bL12 on the ribosome, only a single set of splittings could be resolved, indicating that the conformations and alignment of the CTDs were effectively averaged over the ms chemical-shift timescale. RDCs in the CTD of isolated bL12 (bL12_free_) were of a similar magnitude to those of the domain on the ribosome, but alignment of the isolated NTD (whose resonances are unobservable in the ribosome-bound form) appeared to be significantly weaker, with RDC values of only a few Hz (Figure 1b). These small RDCs are probably due to weaker interactions between the NTD and the phage (both negatively charged)^21^, leading to correspondingly weaker alignment relative to the CTD.

The measured RDCs for the CTD were fitted by singular-value-decomposition (SVD) to a template structure (PDB code: 1rqu^22^) to determine the alignment tensor, *A*^23^. The corresponding back-calculated RDCs showed good agreement with observed RDCs for both bL12_ribo_ and bL12_free_ (*Q* factors of 0.30 and 0.25 respectively, Table S1 and Fig. S4), although for both datasets the RMS deviation (RMSD) between measured and back-calculated RDCs was larger than the RMS uncertainty in the measurements (Table S1). Thus, while the structures of both bL12_ribo_ and bL12_free_ CTD are clearly similar to the template structure, some additional information appears to be present. This finding suggests the possibility of using the RDC data to derive improved structures for both bL12_ribo_ and bL12_free_.

As N-H RDCs could only be measured in a single alignment medium, structure refinement was restricted to secondary-structure elements (SI methods). This procedure resulted in small changes for both bL12_ribo_ and bL12_free_ structures (Fig. 1d), with C*α* RMSDs (relative to the template structure) of 0.35 Å and 0.52 Å respectively, while *Q* factors improved slightly to 0.24 and 0.18 respectively (Table S2). Although not dramatic, this calculation is nevertheless an important step towards the use of RDCs for structural refinement in more complex systems, such as disordered RNCs.

Following refinement, we used the resulting structures to re-fit alignment tensors to the observed RDCs and so compare the preferred orientations of free and ribosome-bound bL12. The orientations of the principal axes of the alignment tensors can be visualised in a Sanson-Flamsteed projection (Fig. 1e), where numerical uncertainties were calculated using a Monte Carlo approach^24^. We observe a small but significant difference in the orientations of the *A_xx_* and *A_yy_* axes (corresponding to a 5D angle between the alignment tensor components of 13.8 ± 7.6º^25^). This result indicates that the orientation of the bL12_ribo_ CTD is influenced by the presence of the core ribosome particle, therefore implying that the bL12 linker region is not completely flexible.

To investigate further the properties of the linker region, we introduced a surface mutation, S89C, in the CTD of bL12 in order to incorporate a lanthanide-binding tag (LBT), Tm-DOTA-M8-4R4S^26^ and induce paramagnetic alignment of the tagged domain (Fig. 2a). The other domains within the bL12 dimer will therefore only exhibit alignment if there is propagation of the alignment from the tagged CTD via the linker(s). The S89C mutation was shown not to perturb the structure of the domain or its dimer formation (as judged by analysis of chemical shift perturbations and diffusion measurements, Fig S5). Two samples were prepared using mixed ^15^N/LBT labelling schemes to distinguish between direct and indirect domain alignment (Fig. 2a). The directly tagged CTD showed strong alignment (maximum observed RDCs of ca. 20 Hz, Fig. 2b and Table S3) with RDCs that fitted well to the template structure (*Q* = 0.15). Alignments of the NTD, and of the untagged CTD (separated by two linkers from the LBT-CTD), were weaker but non-zero (Fig. 2b and Table S3), indicating that some alignment is propagated through the linker region.

**Figure 2.**
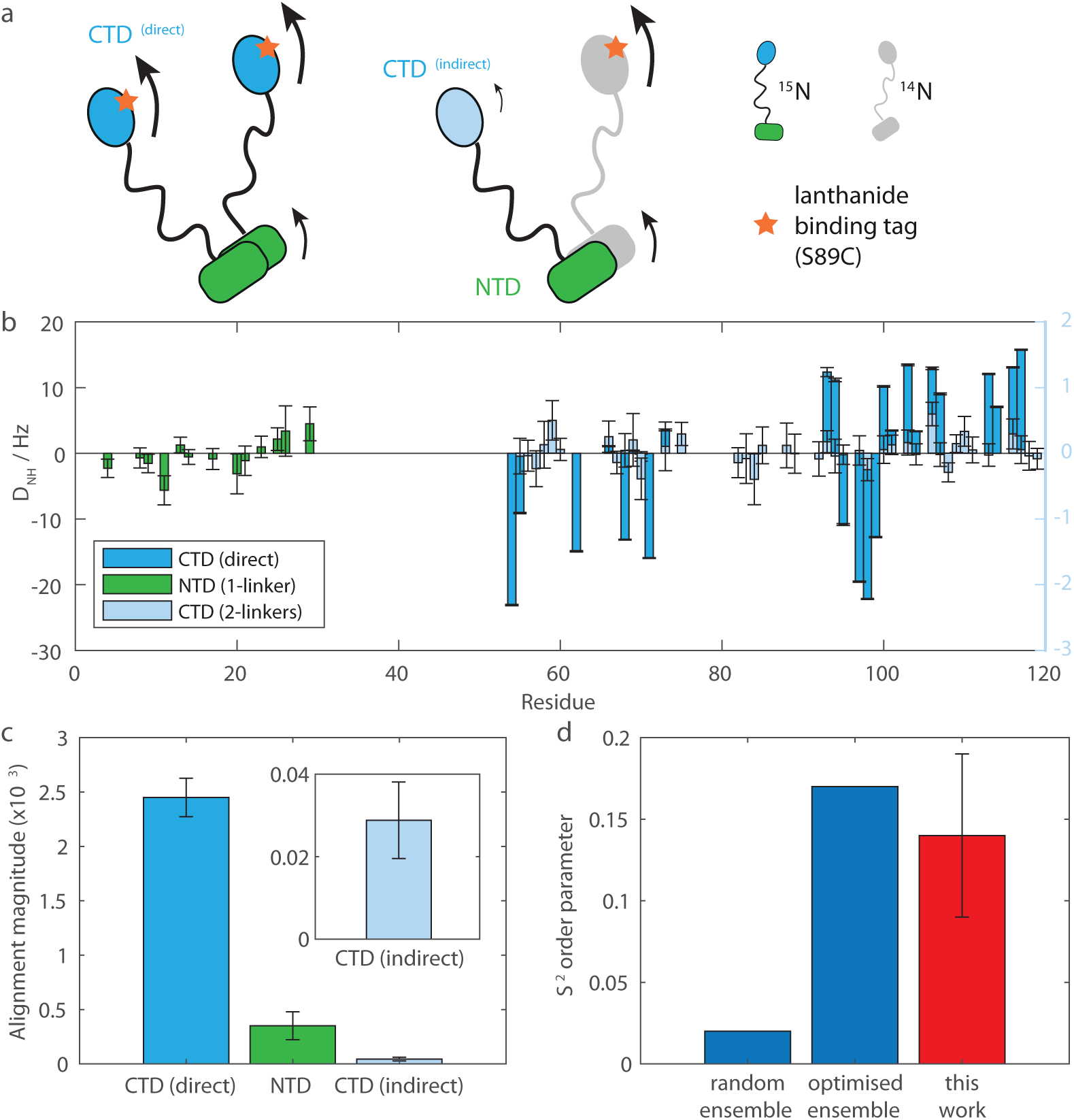
(a) Mixed ^15^N/lanthanide-binding-tag labelling schemes for measurement of the paramagnetic alignment of domains. (b) Amide RDCs measured for directly and indirectly paramagnetically aligned bL12 domains. (c) Magnitude of fitted alignment tensors for directly and indirectly aligned bL12 domains. (d) bL12 linker order parameters determined experimentally (red) and calculated from random and SAXS/NMR-optimised ensembles (blue)^27^.

The alignment strengths of the different domains can be quantified from the magnitude of alignment tensors determined by SVD fitting of the observed RDCs (Fig. 2c). The ratio of alignment magnitudes between adjacent domains then defines an order parameter, *S*^2^ = 0.14 ± 0.05, describing the degree of coupling taking place through the linker region. This measurement can be compared with previous predictions based upon: (i) a random ensemble with a fully flexible linker, and (ii) an optimised ensemble refined using SAXS and ^15^N nuclear spin relaxation measurements (Fig. 2d)^27^. Our measurements are consistent with the latter ensemble, an interesting feature of which is the anti-correlated extension and retraction of the two CTDs, which could potentially assist in the efficient recruitment of elongation factors to the ribosome.

## Conclusions

In summary, we have shown that RDCs can be successfully measured by solution-state NMR spectroscopy for flexible regions of the 2.4 MDa 70S ribosome. The analysis of these measurements indicated a constraining effect of the ribosome on the alignment of the CTDs of the bL12 protein, a conclusion that was verified by independent measurements using specific paramagnetic alignment of domains. Importantly, this study has laid the foundations for detailed characterization of modified bL12 variants on the ribosome and also for more detailed structural investigations into the co-translational folding of ribosome–nascent chain complexes^3, 4^.

## Methods

### NMR Sample preparation

Uniformly ^15^N-labeled ribosomes were prepared by growing *E. coli* cells in MDG^28^ medium containing ^15^NH_4_Cl as the nitrogen source for 20 hours at 310 K before harvesting. In order to escape stationary phase and to ensure that the ribosomes are not hibernating, the cell pellet was then resuspended in an equal volume of fresh medium and incubated at 37 °C until a steady increase of OD_600_ allowed harvesting at growth phase. The resulting cell pellet was lysed by French press in purification buffer (30 mM HEPES, 12 mM Mg(OAc)_2_, 1 M KOAc, 5 mM EDTA, 2 mM beta-mercaptoethanol (BME), pH 7.6) supplemented with trace of DNAse. The soluble fraction of the lysate was purified with a 35 % (w/v) sucrose cushion prepared in purification buffer supplemented with 5 mM ATP. The resulting ribosomal pellet was further purified through a linear density gradient of 10–35 % (w/v) sucrose. Fractions containing 70S ribosomes were identified by SDS-PAGE and pure 70S ribosomes were concentrated and buffer exchanged into Tico buffer (10 mM HEPES, 12 mM MgCl_2_, 30 mM NH_4_Cl, 5 mM EDTA, 2 mM BME, pH 7.6). The concentration of ribosomes was determined from the absorbance at 260 nm using the extinction coefficient *ε* = 4.2 × 10^7^ M^−1^ cm^−120^.

Uniformly ^15^N-labeled/unlabeled bL12 proteins were prepared by transformation into *E. coli* cells and growth in M9 minimal medium containing ^15^NH_4_Cl as the nitrogen source at 37 °C. When the OD_600_ reading reached 0.6, expression was induced with 1 mM IPTG for 4 h at 37 °C. The harvested cell pellet was lysed and purified with a sucrose cushion as described above. bL12 protein was purified from the supernatant using a Hi-Trap Q HP column (loaded in 50 mM Tris-HCl, pH 8.0; elution with a linear gradient of 0–1 M NaCl), followed by a Hi-Trap Phenyl column (loaded in 25 mM Na_2_HPO_4_, 1.7 M (NH_4_)_2_SO_4_, pH 7.5; elution with a reverse gradient of 1.7–1 M (NH_4_)_2_SO_4_) and finally size exclusion chromatography with an Superdex-200 column (50 mM Na_2_HPO_4_, 100 mM KCl, pH 7.0). The concentration of bL12 protein was determined using the bicinchoninic acid (BCA) protein assay reagent (Pierce).

NMR sample concentrations used were 10 *µ*M for the ribosomal particle and 50 *µ*M for isolated bL12. To measure RDCs on the 70S ribosome particle, aligned 70S ribosome samples were prepared by addition of 15 mg mL^−1^ Pf1 filamentous phage (ASLA). For isolated bL12, alignment was achieved by addition of 6.5 mg mL^−1^ filamentous phage. The homogeneity of the resulting anisotropic liquid crystalline medium was confirmed by monitoring the splitting and lineshape of _2_H solvent signal^29^ (Figure S1). The other alignment medium tested (C_12_E_5_ PEG and hexanol^18^) was found not to be suitable due to a strong interaction with isolated bL12 protein. In addition, the presence of ribosomes appeared to interfere with the formation of the homogeneously aligned lamellar phase, as judged from the irregular lineshape of the ^2^H solvent signal (Figure S1C). This latter effect is most likely to arise from the diameter of the ribosomal particle being larger than the inter-lamellar spacing at the PEG concentration used.

To induce alignment using the Tm-DOTA-M8-4R4S^26^ lanthanide binding tag (LBT), cysteine mutations were introduced at positions S15, S24 and S89. All variants were analysed by 2D NMR to confirm that there were no major perturbations of the structure. However, LBT attachment was found by NMR diffusion measurements to disrupt dimer formation in the S15C and S24C variants and therefore only the S89C variant was used for further RDC measurements. For labeling, bL12 (S15C, S24C and S89C) protein was incubated in buffer containing 10 mM DTT and then buffer exchanged into a non-DTT containing buffer on a PD-10 desalting column and reacted with the LBT immediately. The reaction was followed by ^1^H,^1^5N SOFAST-HMQC NMR at 298 K until there was no further change in the spectrum.

### NMR data acquisition

All NMR experiments were recorded at 298K on a Bruker Avance III 700 MHz spectrometer equipped with a TCI cryoprobe. Data were acquired by recording a repeated series of relatively short (time) experiments in an interleaved manner. This approach allows the use of 1D and diffusion experiments to monitor the ribosome integrity at regular intervals. The final spectra for measurement of coupling constants were obtained by summing the individual experiments over the time period for which the ribosome was deemed to be intact, as assessed by ^1^H STE diffusion experiments^30^. These were recorded using an encoding/decoding gradient length, *δ*, of 4.8 ms, a diffusion delay, Δ, of 400 ms, and gradient strengths that were 5, 35, 65 and 95% of the maximum, 55.57 G cm^−1^. To exclude interfering low molecular weight components, the integrity of ribosome samples was monitored using the intensity ratio between 65% and 95% gradient strengths (Figure S3). An initial decrease in the diffusion coefficient for the isotropic sample (Figure S3) can be attributed to a poor baseline due to the residual water signal, as the diffusion coefficients calculated from experiments later in time drops back to ca. 2 × 10^−11^ m^2^s^−1^, which corresponds to intact 70S ribosome^20^. The slower diffusion observed for the aligned sample is a result of elevated viscosity due to the presence of phage (Figure S3B).

Both in-phase/anti-phase (IPAP) ^1^H-coupled ^15^N-HSQC spectra and the pairwise combination of ^15^N-HSQC/^15^N-TROSY spectra were recorded, and the combination of ^15^N-HSQC/^15^N-TROSY spectra was found to yield slightly smaller relative uncertainties in the calculated RDCs than with the approach where two sub-spectra containing the downfield and upfield lines of the ^15^N doublets are generated by taking the sum and difference between in-phase and anti-phase _1_H-coupled ^15^N-HSQC spectra (Figure S2).

### RDC data analysis

Residual dipolar couplings, *D*, were measured as the difference between the isotropic and anisotropic splittings in the ^15^N dimension of HSQC and TROSY spectra. Uncertainties, *σ_D_*, were propagated using standard methods from the uncertainty in peak positions, determined from the signal-to-noise ratio (SN) and ^15^N linewidth (LW), *σ*_ν_ = LW/2SN. Initial data analysis was performed by fitting the measured RDCs to the available NMR-derived structure of the bL12 protein (PDB code 1rqu^22^), as implemented by the SVD (singular-value-decomposition) routine in the program PALES^23^, in order to derive the alignment tensor (*A*) and the corresponding set of back-calculated RDCs. To be able to restrict the analysis of RDCs to the structure and orientational probability distribution of the domain, but not the local dynamics within the domain, residues with local flexibility (order parameter *S*^2^ < 0.85) or undergoing conformational exchange are excluded from the structure and alignment tensor analysis. In addition, residues whose resonances are overlapped in the spectra and residues with experimental uncertainties *σ_D_* > 5 Hz are excluded.

### Structure refinement

The template structure for the CTD of bL12 was refined using Xplor-NIH^31^. The refinement was implemented in a simulating-annealing molecular-dynamics simulation by introducing an additional term to the force-field whose energy depends on the difference between the experimental RDC and that back-calculated from the structure. In the refinement, only the orientation of the secondary-structure elements (individual *α*-helices and *β*-strands) is optimised^32^ and the local structure of the secondary structure elements was fixed using non-crystallographic symmetry (NCS) restraints, and only those RDCs corresponding to rigid N-H vectors within the secondary structure elements were included as restraints. The values for the magnitude and rhombicity of the alignment tensor for back-calculation of the RDCs were taken as those of the best-fit alignment tensor derived using the template structure. The orientation of the alignment tensor is allowed to float during the refinement. The RDC restraints were implemented as soft-square-well susceptibility anisotropy (SANI) restraints, with the width of the well adjusted according to the experimental uncertainty (*σ_D_*) of each measured RDC.

## Acknowledgements

We acknowledge the use of the UCL NMR Centre, the MRC for access to the Biomedical NMR Centre at the Francis Crick Institute, London, and the staff for their support. This work was supported by a Wellcome Trust Investigator Award to J.C. and a Swiss National Science Foundation research grant (SNF 200021 130263) to D.H.

## Author contributions statement

X.W., J.P.K., H.M.M.L., C.M.D., M.V., L.D.C., C.A.W. and J.C. conceived the experiments. X.W., J.P.K. and H.M.M.L. conducted the experiments. D.H. provided reagents. A.d.S. and M.V. performed structure calculations. All authors analysed and discussed the results. X.W., C.A.W. and J.C. wrote the manuscript with input from all authors.

## Additional information

Competing financial interests The authors declare no competing interests.

